# Centrosome depletion rewires mitosis to impose dependence on the AURKA–TPX2 axis

**DOI:** 10.64898/2026.05.02.722379

**Authors:** Zhong Y. Yeow, Fang-Chi Chang, Lance Y. Xu, Andrew J. Holland

## Abstract

Centrosomes are key microtubule-organizing centers required for accurate spindle assembly and chromosome segregation, and their dysfunction in cancer creates therapeutic vulnerabilities. Prior work identified a synthetic lethal interaction between TRIM37 overexpression and Polo-like kinase 4 inhibition (PLK4i) in 17q23-amplified tumors, motivating the clinical development of centrosome-depleting PLK4 inhibitors. However, the broader determinants of sensitivity and resistance to PLK4 inhibition remain poorly defined. Using genome-wide CRISPR–Cas9 screening, we identify multiple genetic suppressors of sensitivity to centrosome depletion, including loss of *PPP6C* as a general escape mechanism, mediated by enhanced activation of Aurora kinase A (AURKA) on the spindle. This process requires NuMA, which scaffolds robust acentrosomal spindle assembly, and operates independently of the TRIM37-regulated pathway that restores pericentriolar material (PCM) foci to reconstitute microtubule-organizing center activity. We further show that centrosome depletion creates a dependence on the AURKA–TPX2 axis for spindle assembly, such that modulation of this pathway shapes cellular responses to PLK4 inhibition. Loss of PPP6C elevates AURKA activity and confers resistance, whereas disruption of the AURKA–TPX2 axis sensitizes cells to centrosome depletion. Together, these findings reveal how centrosome depletion rewires mitotic organization, rendering cells dependent on distinct adaptive spindle assembly pathways.

---

Cancer is characterized by uncontrolled cellular proliferation, placing high demands on the fidelity of mitosis. Early treatments, such as microtubule-targeting agents like taxanes and vinca alkaloids, disrupted spindle assembly to induce mitotic errors and cell death^1^. While effective, these agents are limited by their lack of selectivity, killing healthy dividing cells and causing toxicity in sensory neurons that rely on microtubules for axonal transport^2^. To address these challenges, therapeutic strategies have shifted to exploit cancer-specific vulnerabilities, with synthetic lethality emerging as a central framework in precision oncology^3^. Synthetic lethality occurs when the simultaneous disruption of two genes leads to cell death, while perturbation of either gene alone is tolerated^4^. Synthetic interactions are now being explored across multiple pathways, including those governing epigenetic regulation, metabolism, and cell division^5^.

Centrosomes are large, multiprotein microtubule-organizing centers (MTOCs) that orchestrate spindle assembly during cell division, ensuring accurate chromosome segregation and maintaining genomic stability^6^. Centrosome dysfunction is frequently observed in human tumors and is a common feature across many cancer types^7,8^. Experimental depletion of centrosomes via inhibition of the centriole duplication kinase PLK4 provides a tractable experimental system to study how cells adapt to centrosome loss^9^. In the absence of centrosomes, the formation of surrogate MTOCs such as pericentriolar material (PCM) foci supports spindle assembly^10,11^.

Prior work demonstrated that cancers harboring the 17q23 amplicon are particularly vulnerable to centrosome depletion resulting from PLK4 inhibition^12,13^. This vulnerability arises from the amplicon-driven overexpression of TRIM37, an E3 ubiquitin ligase that degrades PCM components, thereby preventing spindle assembly and triggering mitotic catastrophe^13,14^.

In this study, we sought to define TRIM37-independent mechanisms that govern cellular responses to centrosome loss. Using a genome-wide CRISPR–Cas9 screen, we identified the AURKA–TPX2 axis as a key pathway supporting spindle assembly in centrosome-depleted cancer cells. Mechanistically, loss of PPP6C suppresses PLK4i hypersensitivity by enhancing spindle-localized AURKA activity, thereby promoting robust acentrosomal spindle assembly through a pathway distinct from TRIM37-mediated PCM restoration. We further demonstrate that perturbation of the AURKA–TPX2 axis sensitizes cells to PLK4 inhibition. Together, these findings establish a framework in which distinct compensatory pathways are engaged to sustain spindle assembly following centrosome depletion, thereby underpinning cellular responses to PLK4 inhibition.

## Results

### Determinants of cancer cell sensitivity to PLK4 inhibitor–induced centrosome depletion extend beyond TRIM37 overexpression and p53 status

To determine whether sensitivity to centrosome depletion extends beyond known contexts, we screened a panel of cell lines for response to centrinone, a small-molecule PLK4 inhibitor (PLK4i)^9^. Clonogenic survival assays demonstrated that, in addition to the established sensitivity of MCF-7 cells (which harbor 17q23 amplification and overexpress TRIM37)^13^, both HeLa and HCC1806 cells exhibited strong sensitivity to PLK4i treatment (Fig. 1a). Western blot analysis revealed that only MCF-7 cells overexpressed TRIM37, while HeLa and HCC1806 cells did not (Extended Data Fig. 1a). Notably, PLK4i-treated RPE-1 TP53*^-/-^* and DLD1 cells were largely unaffected by PLK4i treatment (Fig. 1a), consistent with their ability to form PCM foci that support spindle assembly in the absence of centrosomes^11,13^.

**Figure 1.**
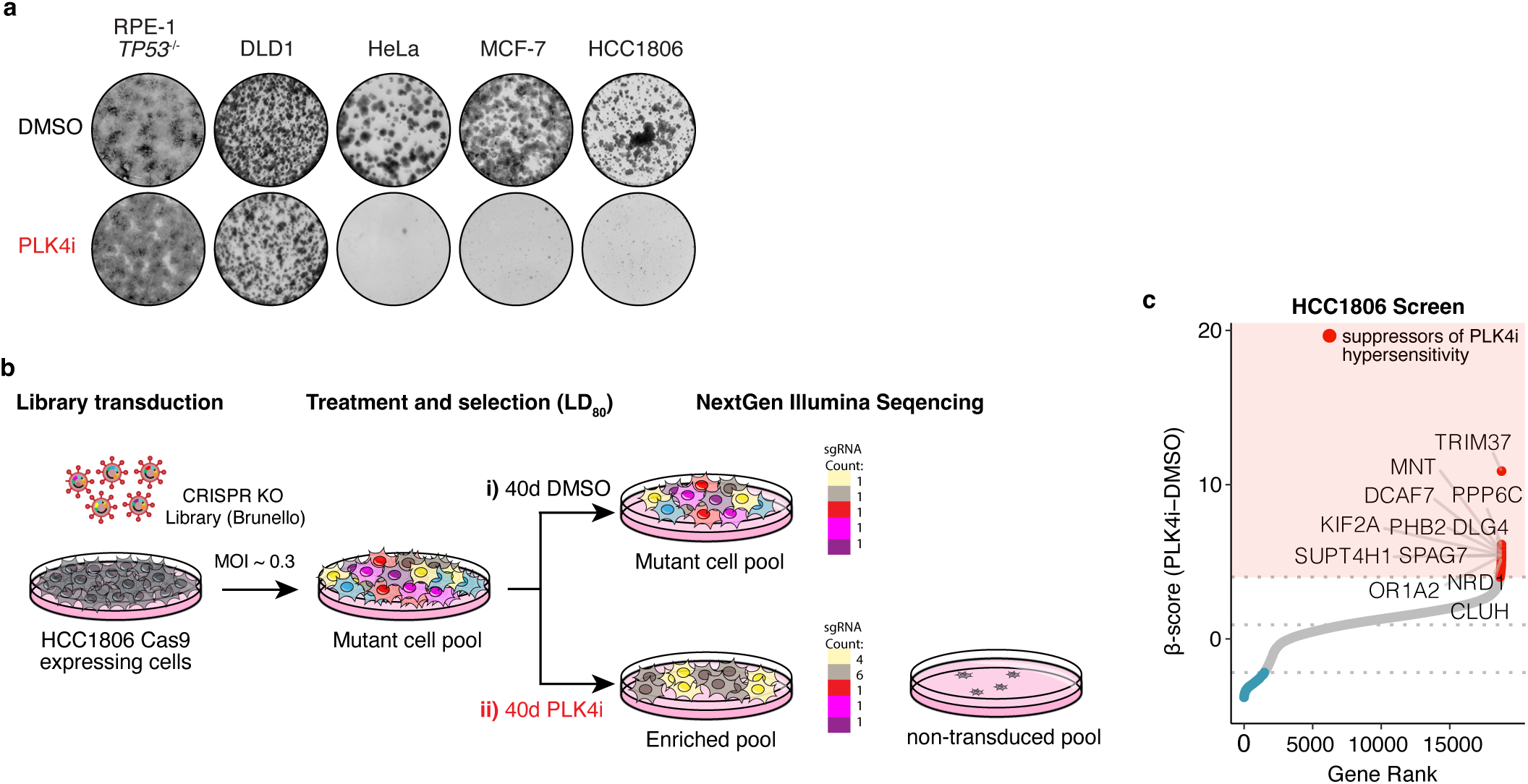
CRISPR–Cas9 screen reveals genetic suppressors of PLK4 inhibitor sensitivity. **a**, Representative 14-d clonogenic survival assay of RPE-1 *TP53^-/-^*, DLD1, HeLa, MCF-7 and HCC1806 cell lines treated with DMSO (control) or the PLK4 inhibitor centrinone (PLK4i). *n* = 3 biological replicates. **b**, Schematic of the genome-wide CRISPR–Cas9 knockout screen used to identify genes whose loss suppresses PLK4i hypersensitivity in HCC1806 cells. LD_80_, lethal dose 80. **c**, CRISPR screen results plotted by β-score (PLK4i – DMSO) versus gene rank. The top 12 genes enriched under PLK4i treatment and exceeding 2 × SD above the population mean (grey dotted lines) are highlighted. SD, standard deviation.

Centrosome loss in non-transformed cells triggers the p53-mediated mitotic surveillance pathway^15^, leading to cell cycle arrest. To determine whether HeLa and HCC1806 cells possess a functional p53 response, we treated the cell lines with the p53 activator Nutlin-3a^16^. MCF-7 cells arrested in response to Nutlin-3a treatment, indicating functional p53 (Extended Data Fig. 1b). However, HeLa and HCC1806 cells continued to proliferate, showing that the p53 pathway is defective in these cells. Together, our data show that the sensitivity of HeLa and HCC1806 cells to PLK4 inhibition is independent of both TRIM37 overexpression and p53-mediated growth arrest, highlighting a dependence on additional pathways following centrosome depletion.

### CRISPR screen identifies suppressors of sensitivity to centrosome depletion

To understand the mechanisms governing cellular responses to centrosome depletion induced by PLK4 inhibition, we performed a genome-wide CRISPR–Cas9 knockout screen in HCC1806 Cas9-expressing cells. The human Brunello sgRNA library was transduced and cells treated with DMSO or PLK4i for 40 days before next-generation sequencing^17^ (Fig. 1b). Since PLK4 inhibition depletes centrosomes, our screen was designed to enrich for genes whose loss rescues acentrosomal spindle assembly, thus conferring resistance to PLK4i (Extended Data Fig. 1c). Four genes with established roles in mitosis emerged as top hits (Fig. 1c): *TRIM37*^12,13,18–20^, *MNT*^21,22^, *KIF2A*^23^, and *PPP6C*^24,25^.

While TRIM37 is not overexpressed in HCC1806 cells (Extended Data Fig. 1a), its knockout suppressed hypersensitivity to PLK4 inhibition (Extended Data Fig. 2a, b). This suppression was associated with the formation of acentrosomal PCM foci marked by CEP192 (Extended Data Fig. 2c, d) and is consistent with prior work showing that TRIM37 loss accelerates acentrosomal spindle assembly following centrosome depletion^13^. Spindle analysis showed that TRIM37 knockout restored bipolar spindle formation and normal mitotic progression in HCC1806 cells following PLK4i-induced centrosome depletion (Extended Data Fig. 2c, e, f). These findings show that although HCC1806 cells do not overexpress TRIM37, TRIM37 knockout nevertheless restores PCM foci formation to support acentrosomal spindle assembly and rescues cell growth under PLK4i treatment^12,13^.

### *PPP6C* knockdown rescues acentrosomal mitosis in PLK4i-treated cancer cells

Having validated the screening hit *TRIM37*, we next focused on *PPP6C*, another top-ranked hit (Fig. 1c). *PPP6C* encodes for the catalytic subunit of protein phosphatase 6 (PP6), a negative regulator of Aurora kinase A (AURKA)^25^. To investigate how *PPP6C* disruption impacts PLK4i sensitivity, we generated two monoclonal *PPP6C*-edited (hereafter sg*PPP6C*) cell lines. PPP6C knockdown was confirmed by immunoblotting and resulted in a marked suppression of PLK4i hypersensitivity compared to sgEV control (Fig. 2a, b). Upon centrosome depletion, bipolar spindle formation was significantly impaired in PLK4i-treated sgEV cells, with only 44.6% of cells forming bipolar spindles (Extended Data Fig. 3a–c). This was accompanied by an increased incidence of monopolar (19.5%), disorganized (26.7%), and multipolar spindles (9.2%). These spindle defects were markedly reduced in sg*PPP6C* cells, with 75.8% of PLK4i-treated cells assembling bipolar spindles (Extended Data Fig. 3a–c). Notably, this rescue occurred without the formation of detectable PCM (CEP192) foci in mitosis, indicating a distinct rescue pathway from loss of TRIM37 (Extended Data Fig. 2c, Extended Data Fig. 3a). Additionally, mitotic phase analysis showed that PPP6C knockdown reduced the prometaphase population from 58% in sgEV controls to 25% in sg*PPP6C* cells (Extended Data Fig. 3a, d), indicating restored progression through mitosis.

**Figure 2.**
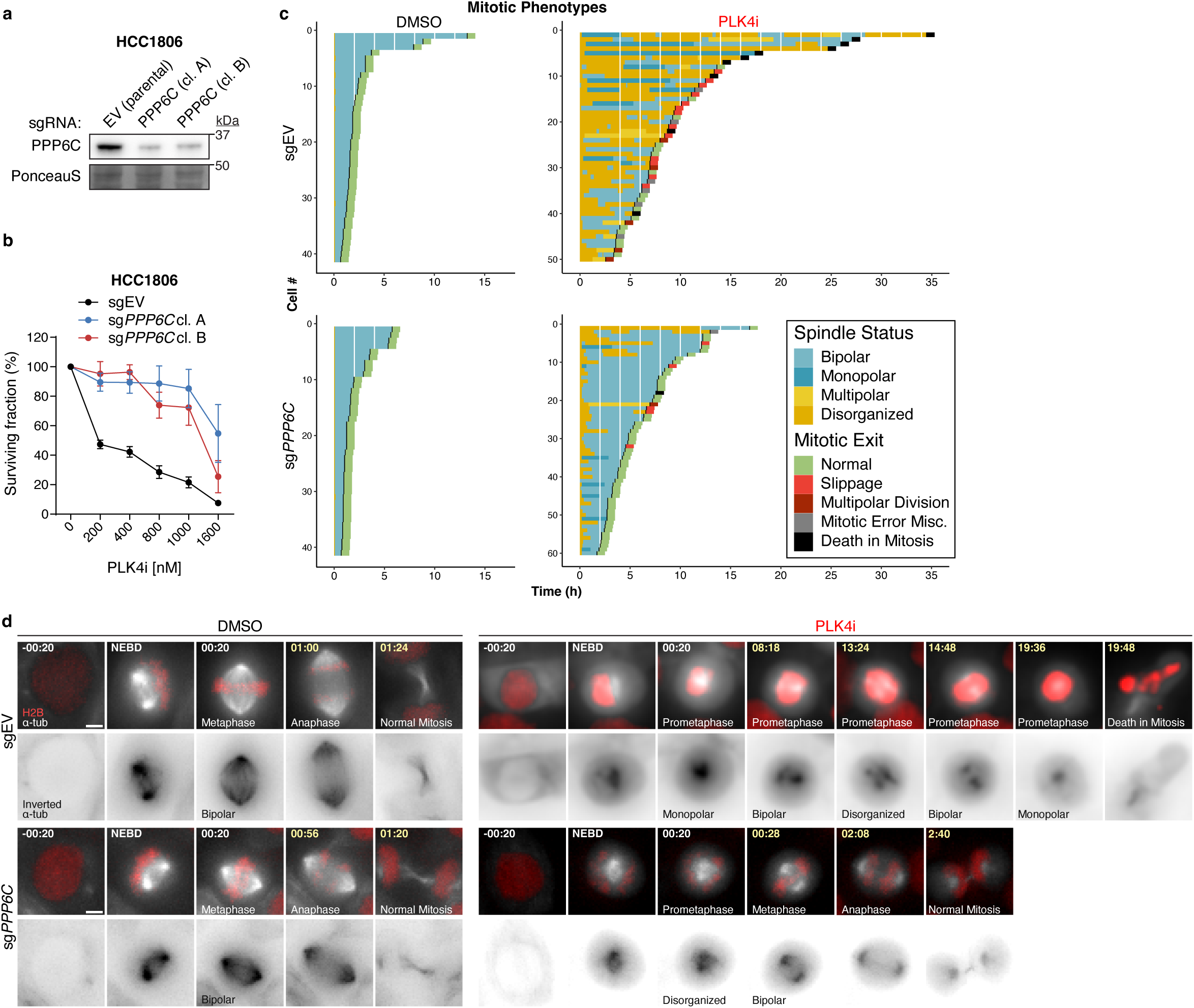
*PPP6C* loss alleviates PLK4i-induced mitotic defects in HCC1806 cells. **a**, Immunoblot showing PPP6C levels in parental HCC1806 and two *PPP6C*-edited clones. Ponceau-stained blot indicates loading. Representative data; *n* = 3 biological replicates. EV, empty vector. **b**, PLK4i sensitivity of the cell lines shown in (**a**) plotted as surviving fraction across increasing PLK4i concentrations and quantified by a resazurin-based viability assay. **c**, Quantification of mitotic duration, spindle configuration, and mitotic exit outcome from live-cell time-lapse imaging of HCC1806 cells treated with DMSO or PLK4i. n > 40 cells per condition across *n* = 3 biological replicates. **d**, Representative time-lapse sequences corresponding to datasets quantified in (**c**), showing mitotic progression of fluorescently tagged H2B–iRFP and EGFP–α-tubulin HCC1806 cells treated with DMSO or PLK4i. Scale bar = 5 μm. NEBD, nuclear envelope breakdown.

To validate these findings, we performed time-lapse imaging to track spindle dynamics and mitotic outcomes. DMSO-treated cells assembled bipolar spindles and progressed through mitosis normally in a timely manner (Fig. 2c, d and Supplementary Videos 1, 3). However, in PLK4i-treated sgEV cells, transient bipolar spindles frequently collapsed into disorganized or monopolar configurations, resulting in prolonged mitoses and adverse outcomes such as mitotic death or slippage (Fig. 2c, d and Supplementary Video 2). In contrast, sg*PPP6C* cells maintained stable bipolar spindles throughout cell division and exited mitosis normally despite PLK4i treatment (Fig. 2c, d and Supplementary Video 4). We conclude that PPP6C knockdown enables robust acentrosomal spindle assembly and restores normal mitotic progression, via a mechanism independent of PCM foci.

### Potentiation of Aurora kinase A (AURKA) activity rescues acentrosomal mitosis and suppresses PLK4i hypersensitivity

PPP6C negatively regulates AURKA by dephosphorylating it at Thr288 when complexed with TPX2^25^. Recognizing this role, we hypothesized that PPP6C knockdown would enhance AURKA activity, thereby restoring growth in PLK4i. This was supported by the enrichment of AURKA and its active phosphorylated form (AURKA-pT288) at the spindle in sg*PPP6C* cells (Fig. 3a, b), consistent with prior reports^24,25^. Moreover, AURKAi treatment resulted in the loss of centrosomal AURKA levels, whereas PPP6C knockdown partially restored the level of AURKA at centrosomes (Fig. 3a). The loss of PPP6C also substantially increased the resistance of HCC1806 cells to the AURKA inhibitor MLN8237 (Fig. 3c), supporting the proposal that AURKA activity is elevated in PPP6C knockout cells.

**Figure 3.**
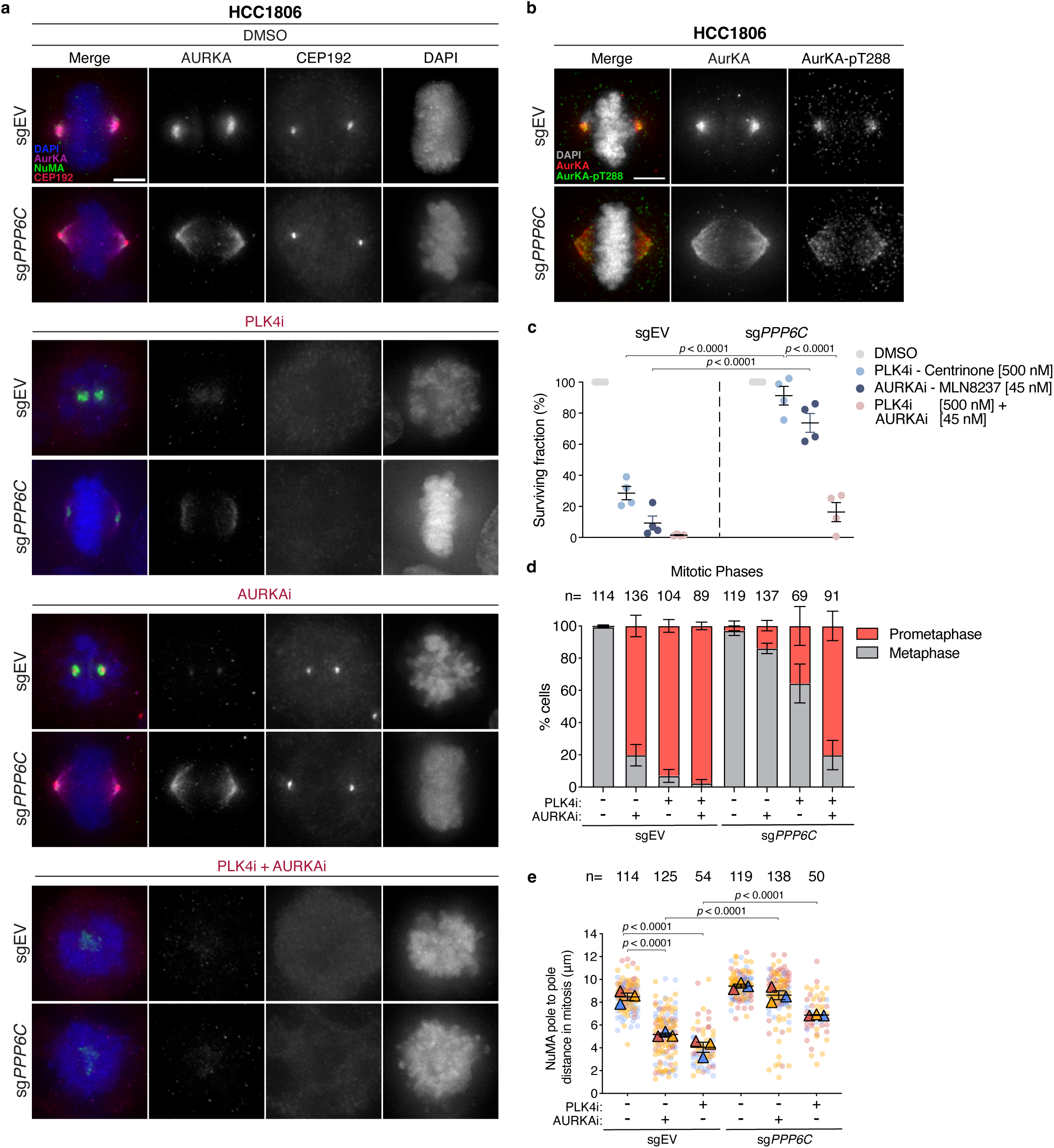
PPP6C loss potentiates AURKA activity to restore acentrosomal mitosis under PLK4 inhibition. **a**, Representative immunofluorescence images of mitotic HCC1806 sgEV and sg*PPP6C* cells treated with DMSO (control), PLK4i (500 nM), AURKAi (45 nM), or the combination of PLK4i and AURKAi for 6 d. *n* = 3 biological replicates. Scale bar = 5 μm. **b,** Representative immunofluorescence images of mitotic HCC1806 sgEV and sg*PPP6C* cells stained for total AURKA and its active, Thr288-phosphorylated form (pT288). **c**, Surviving fraction of HCC1806 sgEV and sg*PPP6C* cells treated under the conditions shown in (**a**) for 10 d, quantified by a resazurin-based viability assay. *n* = 4 biological replicates. *P* values were determined using a two-way ANOVA with Sidak’s multiple-comparisons test. Mean ± s.e.m. **d**, Quantification of the percentage of cells in each mitotic phase under the treatment conditions shown in (**a**). *n* = 3 biological replicates. **e**, Quantification of pole-to-pole distance in mitotic cells under the treatment conditions shown in (**a**). Triangles represent the mean for each biological replicate; coloured circles show individual data points from each of the replicates. *n* = 3 biological replicates. *P* values were determined using a two-way ANOVA with Sidak’s multiple-comparisons test. Mean ± s.e.m.

We reasoned that if sg*PPP6C*-mediated suppression of PLK4i sensitivity relies on enhanced AURKA activity, then inhibiting AURKA should re-sensitize these cells to PLK4i. Consistent with this hypothesis, AURKA inhibition rendered sg*PPP6C* cells sensitive to PLK4i, as evidenced by a reduction in survival rates to levels comparable to PLK4i-treated sgEV cells (Fig. 3c). This re-sensitization was accompanied by an increase in prometaphase-arrested cells, replicating the phenotype observed with PLK4i alone in sgEV cells (Fig. 3a, d). Together, these findings demonstrate that PPP6C knockdown restores acentrosomal spindle assembly and mitotic progression by potentiating AURKA activity.

### NuMA acts as a scaffold that supports acentrosomal spindle assembly

During acentrosomal cell division, pericentriolar material (PCM) proteins such as CEP192 typically form microtubule-organizing centers (MTOCs) that facilitate spindle assembly^10,11,13^. Interestingly, in HCC1806 sg*PPP6C* cells, acentrosomal mitosis was rescued in the absence of CEP192 foci (Fig. 3a, Extended Data Fig. 3a, b), suggesting the involvement of alternative scaffolding proteins. Previous studies showed that in the absence of PCM the nuclear mitotic apparatus (NuMA) protein can organize microtubule asters to support acentrosomal spindle formation^26,27^. We therefore examined NuMA localization in HCC1806 sgEV and sg*PPP6C* cells and found it to be consistently present at acentrosomal spindle poles, even in the absence of canonical PCM proteins like CEP192 and CDK5RAP2 (Fig. 4a, Extended Data Fig. 4a, b). Expanding this analysis to other cell lines, we observed that while all three of these PCM proteins were detectable at centrioles in DMSO-treated control cells, PCM retention varied widely upon centrosome loss. PLK4i-insensitive cell lines (RPE-1 *TP53*^-/-^ and DLD1) retained appreciable PCM components at the acentrosomal poles, whereas PLK4i-sensitive cell lines (HeLa, MCF-7, and HCC1806) showed lower overall PCM retention. All three PLK4-sensitive cell lines lacked detectable CEP192, while PCNT was absent at the poles of MCF-7 and HeLa cells but detectable in a proportion of HCC1806 cells (Extended Data Fig. 4a, b). In contrast, NuMA was consistently present at the acentrosomal poles in all cell lines tested, irrespective of PLK4i sensitivity. NuMA typically presented as two distinct foci flanking the metaphase plate or, in HeLa cells, a dense column (Extended Data Fig. 4a, b) as previously reported^26^.

**Figure 4.**
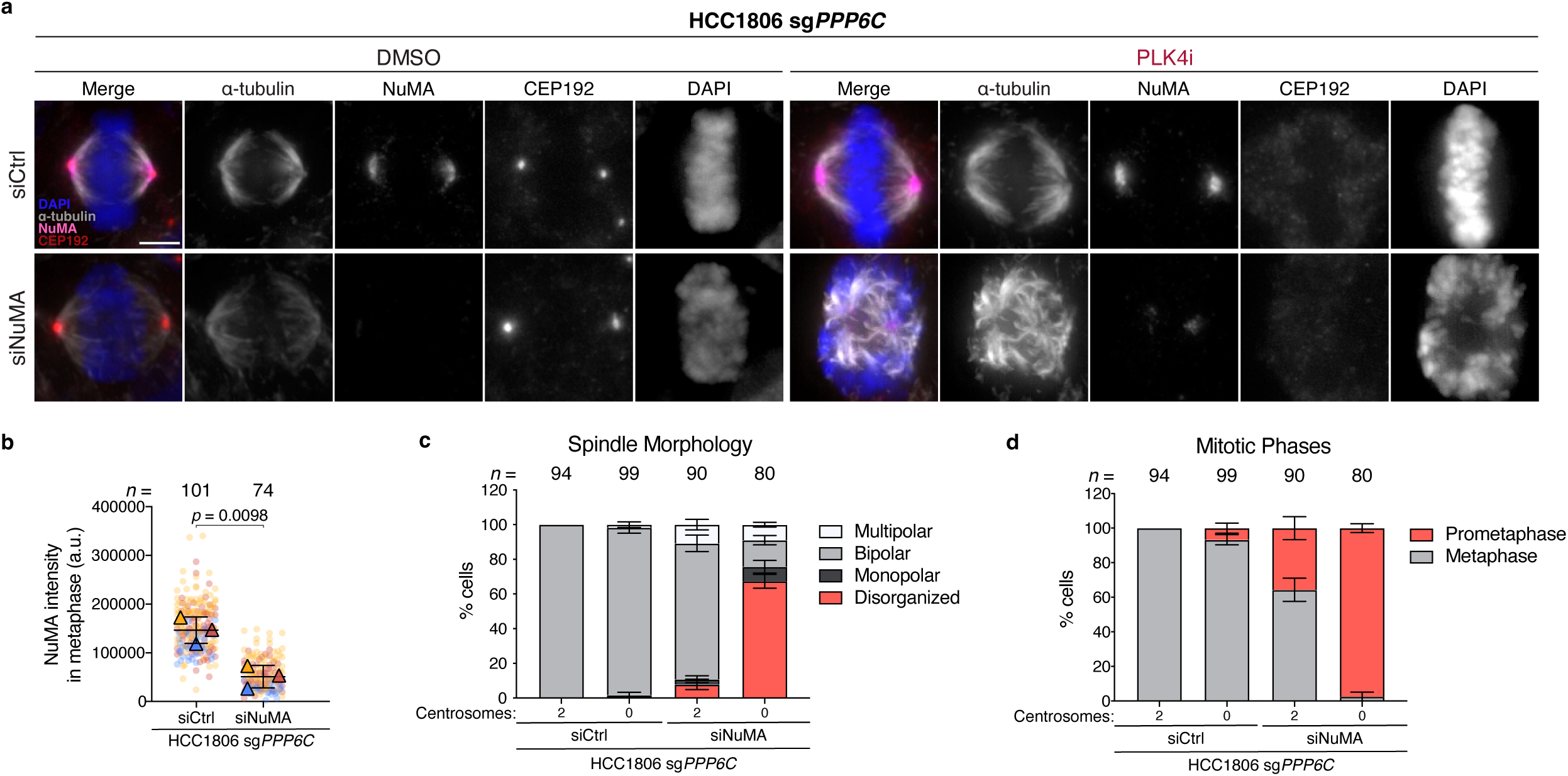
NuMA acts as an essential pole scaffold for acentrosomal spindle assembly in HCC1806 sg*PPP6C* cells. **a**, Representative immunofluorescence images of mitotic HCC1806 sg*PPP6C* cells transfected with non-targeting siCtrl or siNuMA and treated with DMSO (control) or PLK4i (500 nM) for 6 d. Scale bar = 5 μm. Ctrl, control. **b**, Quantification of NuMA intensity in metaphase cells transfected with siCtrl or siNuMA, as shown in (**a)**. Triangles represent the mean for each biological replicate; coloured circles show individual data points from each of the replicates. *n* = 3 biological replicates. *P* values were determined using an unpaired two-tailed *t*-test. Mean ± s.e.m. **c,** Quantification of the percentage of cells displaying the indicated spindle morphology under the treatment conditions shown in (**a**). Mean ± s.e.m. **d,** Quantification of the percentage of cells in each mitotic phase under the treatment conditions shown in (**a**). Mean ± s.e.m.

To dissect NuMA’s role in the acentrosomal mitosis of HCC1806 sg*PPP6C* cells, we performed siRNA-mediated NuMA knockdown and analyzed spindle morphology and mitotic progression (Fig. 4a, b). While NuMA depletion only modestly reduced bipolar spindle assembly in cells with two centrosomes, bipolar spindle formation decreased from 97% to 15% in acentrosomal cells following NuMA depletion (Fig. 4a, c). Additionally, NuMA knockdown led to a significant increase in the prometaphase population compared to control siRNA treatment (97% vs. 7%) (Fig. 4a, d), underscoring NuMA’s role as a scaffold for acentrosomal spindles in the absence of centrosomes and PCM foci. Thus, our results demonstrate that sg*PPP6C*-mediated potentiation of AURKA at the spindle is essential during acentrosomal mitosis, with NuMA acting as a critical scaffold for spindle assembly.

### Centrosome depletion reveals dependence on the AURKA–TPX2 axis for spindle assembly

Previous studies established that the AURKA–TPX2 axis is crucial for setting and maintaining spindle length^28^. In line with this, treatment with AURKAi significantly reduced spindle length in HCC1806 sgEV cells, and this was rescued by hyperactivating AURKA with PPP6C knockdown (Fig. 3e). Since centrosomes act as key hubs for AURKA activation, we reasoned that their loss would reduce AURKA activity, leading to shorter spindles. Accordingly, centrosome loss induced by PLK4i treatment reduced spindle length to a similar extent as AURKA inhibition, and elevating AURKA activity with PPP6C knockdown again restored normal spindle length (Fig. 3e).

To examine how centrosome depletion impacts AURKA activity and spindle architecture, we used short-term PLK4 inhibition to generate cells containing a single centrosome to differentiate centrosome-dependent versus acentrosomal AURKA activity within the same cell. Acentrosomal poles (marked by the absence of CEP192 and presence of NuMA) exhibited lower levels of AURKA and were positioned closer to the metaphase plate than centrosome-containing poles (marked by CEP192) (Extended Data Fig. 5a–c). Knockdown of PPP6C modestly increased the level of AURKA activity at acentrosomal poles, accompanied by an increase in half-spindle length (3.7 μm in sg*PPP6C* cells versus 3.09 μm in sgEV cells) (Extended Data Fig. 5a–c). These observations suggest that PPP6C knockdown enhances AURKA activity at acentrosomal spindles, compensating for the absence of centrosomes.

In PPP6C knockdown cells, enhanced AURKA localization on the spindle stems from its increased association with spindle-localized TPX2^25^. To test whether TPX2 is required for PLK4i resistance in cells lacking PPP6C, we used shRNA to deplete TPX2 and displace AURKA from the spindle (Extended Data Fig. 6a). Loss of TPX2 decreased spindle length^28^ and sensitized cells to PLK4i (Extended Data Fig. 6b, c). These findings underscore that PLK4i resistance in HCC1806 sg*PPP6C* cells is dependent on the spindle-associated pool of active AURKA.

We postulated that HCC1806 cells might harbor an inherent dysfunction in the AURKA–TPX2 axis that is revealed when centrosomes are depleted via PLK4 inhibition. This predicts a heightened sensitivity of HCC1806 cells to AURKA inhibition. To explore this hypothesis, we assessed the sensitivity of HCC1806 cells to AURKA inhibition relative to other cell lines. Growth assays revealed that PLK4i-insensitive cell lines (RPE-1 and DLD-1) were highly resistant to AURKA inhibition. In contrast, HCC1806 cells displayed profound sensitivity to AURKA inhibition, while the other PLK4i-sensitive cell lines (HeLa and MCF-7) exhibited modest sensitivity to AURKAi (Extended Data Fig. 7a). This supports the notion that HCC1806 cells possess a distinct defect in the AURKA–TPX2 axis driving heightened sensitivity to both PLK4 and AURKA inhibition.

To determine whether AURKA dependency can be induced in cell lines without intrinsic AURKA–TPX2 dysfunction, we tested dual inhibition of PLK4 and AURKA in RPE-1 and DLD-1 cells. This combined treatment sensitized these cell lines to PLK4i (Extended Data Figure 7b), indicating that centrosome depletion induces a conditional dependence on AURKA to support spindle assembly.

### Broad suppression of PLK4i sensitivity across cancer cells by PPP6C knockdown

Our findings show that HeLa and MCF-7 cells, like HCC1806, are hypersensitive to PLK4 inhibition (Fig. 1a). To evaluate the broader suppressive effect of PPP6C, we generated sg*PPP6C* lines for both MCF-7 and HeLa. PPP6C knockdown was confirmed by immunoblotting and resulted in increased AURKA localization to the spindle, as well as robust suppression of PLK4i hypersensitivity in both cell lines, mirroring the effects observed in HCC1806 sg*PPP6C* cells (Fig. 5a–c). Following PLK4i-induced centrosome depletion, bipolar spindle formation was substantially impaired in sgEV MCF-7 and HeLa cells (Fig. 5d–f, h–j). Notably, PPP6C knockdown alleviated these spindle defects, resulting in a higher proportion of bipolar spindles under PLK4i treatment (Fig. 5d–f, h–j). Mitotic phase analysis also revealed a marked reduction in prometaphase arrest in PLK4i-treated sg*PPP6C* cells compared to sgEV controls, indicating improved mitotic progression (Fig. 5d, g, h, k). Overall, these results demonstrate that PPP6C knockdown effectively restores acentrosomal spindle assembly, mitigating PLK4i-induced mitotic defects across multiple sensitive cell lines. These findings highlight increased AURKA activity as a general mechanism supporting acentrosomal spindle assembly in the context of centrosome depletion (Fig. 6).

**Figure 5.**
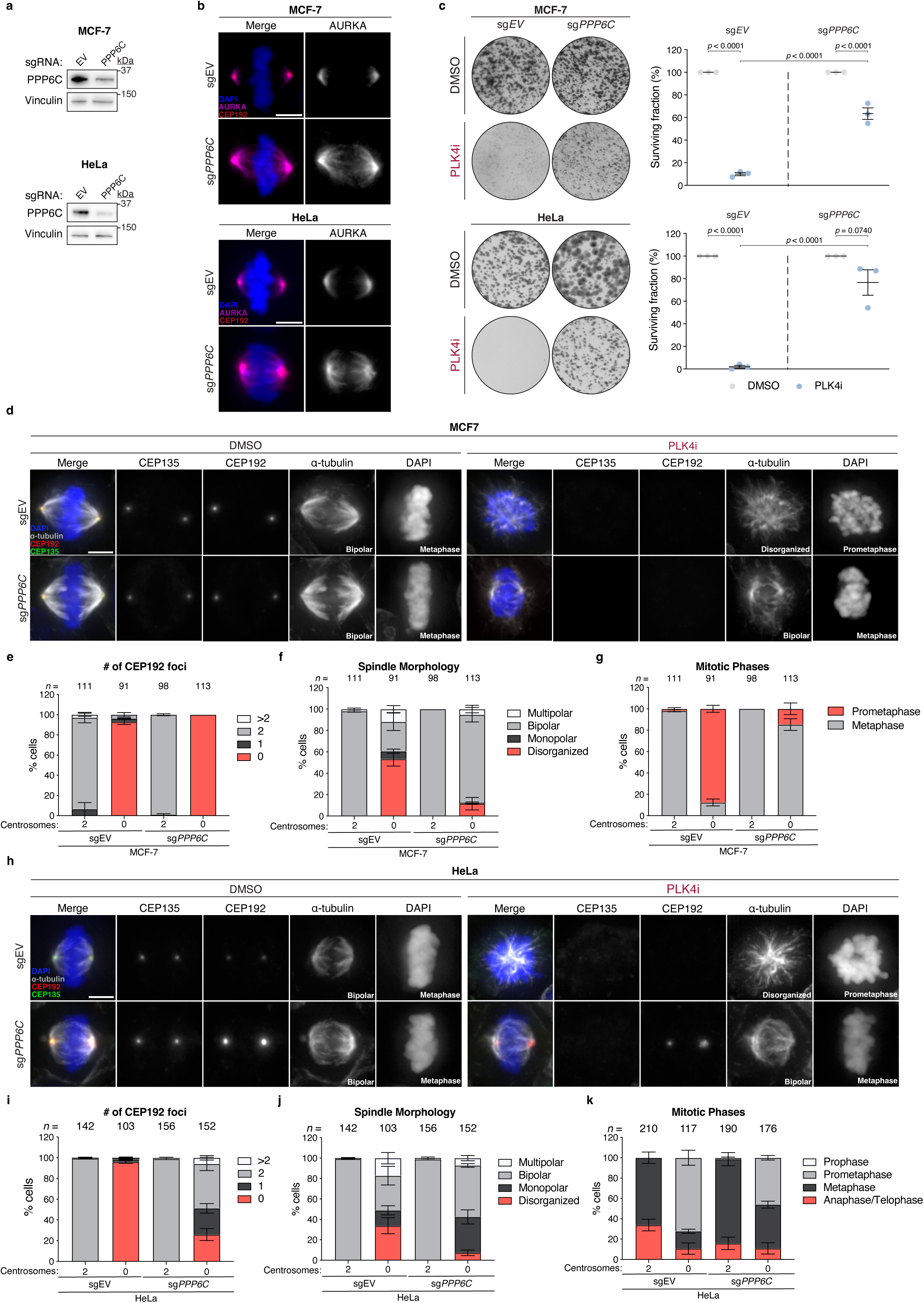
*PPP6C* loss broadly suppresses PLK4i hypersensitivity across multiple cancer cell lines. **a**, Immunoblot showing PPP6C levels in parental MCF-7 and HeLa and their corresponding sg*PPP6C* clones. Vinculin, loading control. Representative data; *n* = 3 biological replicates. EV, empty vector. **b**, Representative immunofluorescence images of metaphase MCF-7 and HeLa cells showing increased spindle-associated AURKA in sg*PPP6C* clones compared to sgEV controls. Scale bar = 5 μm. **c**, Left, Representative 14-d clonogenic survival assay of MCF-7 and HeLa cell lines treated with DMSO (control) or the PLK4 inhibitor centrinone (PLK4i). *n* = 3 biological replicates. Right, Quantification of surviving fraction. *P* values were determined using a one-way ANOVA with a post hoc Tukey’s multiple-comparisons test. Mean ± s.e.m. **d**, Representative immunofluorescence images of mitotic MCF-7 sgEV and sg*PPP6C* cells treated with DMSO (control) or PLK4i (500 nM) for 6 d. Scale bar = 5 μm. **e**–**g**, Quantification of centrosome number (**e**), spindle morphology (**f**), and mitotic phase distribution (**g**) of MCF-7 sgEV and sgPPP6C cells under the treatment conditions shown in **d**. n = 3 biological replicates. Mean ± s.e.m. **h**–**k**, Same analyses as in **(d**–**g)**, performed in HeLa sgEV and sg*PPP6C* cells.

**Figure 6.**
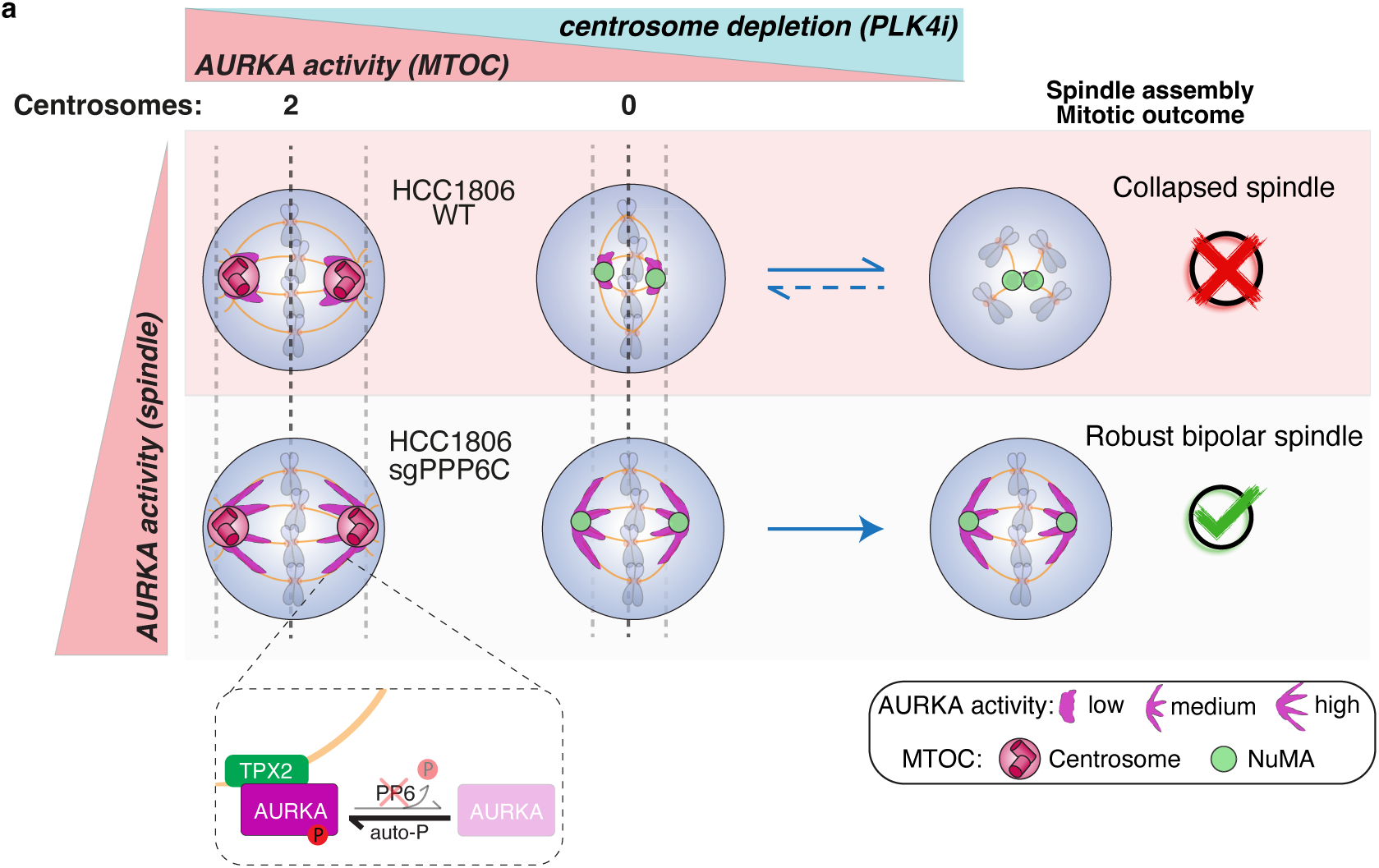
Model illustrating how AURKA activity stabilizes the acentrosomal spindle following centrosome depletion. **a**, Centrosome loss removes the major platform for AURKA activation, leading to reduced AURKA activity at spindle poles and shortened spindles that are prone to collapse. This spindle instability underlies PLK4i hypersensitivity in parental HCC1806 cells. Loss of PPP6C (a negative AURKA regulator) potentiates spindle-associated AURKA activity, enabling cells to maintain robust NuMA-scaffolded acentrosomal spindles and achieve faithful mitotic progression.

## Discussion

Synthetic lethality offers a powerful framework for targeting cancer-specific vulnerabilities^5^. Building on the discovery of synthetic lethality between TRIM37 overexpression and PLK4 inhibition in 17q23-amplified cancers^12,13^, our study identifies additional determinants of sensitivity to centrosome depletion that extend beyond TRIM37-driven contexts. Using a genome-wide CRISPR screen, we uncovered two mechanistically distinct pathways that alleviate PLK4i hypersensitivity. TRIM37 depletion restores spindle assembly by promoting ectopic PCM foci that function as alternative MTOCs, while PPP6C knockdown activates a spindle-centric mechanism that enhances AURKA activity independent of PCM. Together, these findings reveal the remarkable adaptability of acentrosomal mitosis and highlight potential avenues to target these compensatory pathways.

A key insight from our work is the requirement for AURKA and TPX2 to sustain bipolar spindle assembly in the absence of centrosomes. Under normal conditions, AURKA is recruited to centrosomes via PCM proteins such as PCNT and CEP192, where its local concentration promotes activation^29^. PLK4 inhibition depletes centrosomes, reducing AURKA abundance and activity at spindle poles, leading to shortened, fragile spindles. As a consequence, acentrosomal cells become functionally dependent on the TPX2-bound pool of AURKA localized to spindle microtubules, which locally activates AURKA independent of PCM. Hence, the AURKA–TPX2 axis acts as a key mediator of spindle organization and mitotic progression under acentrosomal conditions (Fig. 6). We further demonstrate that this pathway is compromised in HCC1806 cells, which exhibit hypersensitivity to AURKA inhibition at nanomolar concentrations. Importantly, their vulnerability to centrosome depletion is independent of TRIM37 status or p53 function, indicating that alterations in the AURKA–TPX2 axis represent an additional determinant of this response.

Our results also establish NuMA as a critical scaffold for acentrosomal spindle poles, compensating for the absence of canonical PCM components including CEP192, PCNT, and CDK5RAP2. Prior studies highlighted NuMA’s role in organizing acentrosomal asters^26,27^, and we now demonstrate its essentiality in centrosome-depleted HCC1806 cells. CEP192 is known to prime AURKA by generating a pool of T-loop phosphorylated kinase necessary for TPX2 binding and spindle recruitment during mitosis^29^. In the absence of centrosomes, our findings suggest that PPP6C knockdown bypasses this CEP192 dependency by preventing AURKA dephosphorylation on the spindle, thereby elevating AURKA activity and enabling robust bipolar spindle assembly. This plasticity of acentrosomal mitotic pathways parallels natural processes such as oocyte meiosis, where AURKA–TPX2 and NuMA are indispensable^30,31^, underscoring the evolutionary conservation of acentrosomal division mechanisms.

From a therapeutic perspective, our findings illuminate potential resistance mechanisms to PLK4 inhibition. PLK4 inhibitors are currently under clinical evaluation for tumors with high TRIM37 expression (NCT06232408), highlighting the need to understand both determinants of response and resistance in this setting. In 17q23-amplified cancers, genomic amplification drives TRIM37 overexpression, rendering tumors hypersensitive to PLK4i. Our data suggest that AURKA potentiation may represent an adaptive escape mechanism by which cancer cells overcome PLK4i sensitivity. Indeed, PPP6C downregulation in 17q23-amplified MCF-7 cells broadly suppresses PLK4i sensitivity by enhancing spindle-localized AURKA activity and restoring acentrosomal spindle assembly (Fig. 5). AURKA has long been recognized as an oncogenic driver, frequently amplified across various cancer types and associated with clinical aggressiveness and poor prognosis^32,33^. However, clinical use of AURKA inhibitors has been limited by dose-limiting toxicity^32^. Our findings suggest that PLK4i-induced centrosome depletion increases dependency on AURKA, thereby sensitizing cells to AURKA inhibition at reduced doses. Together, these observations raise the possibility that co-targeting AURKA may improve the durability of responses to PLK4i therapy.

## Acknowledgements

This work was supported by research grants R01GM114119, R01GM133897 and R01CA266199 (to A.J.H) from the National Institutes of Health.

## Author contributions

F.C.C and Z.Y.Y. designed, performed, and analysed the majority of the experiments and prepared the figures. L.Y.X assisted with cloning and immunofluorescence analysis. Z.Y.Y. and A.J.H. conceived and supervised the study. Z.Y.Y. and A.J.H. co-wrote the manuscript.

## Extended Data Figure Legends

**Extended Data Figure 1.**
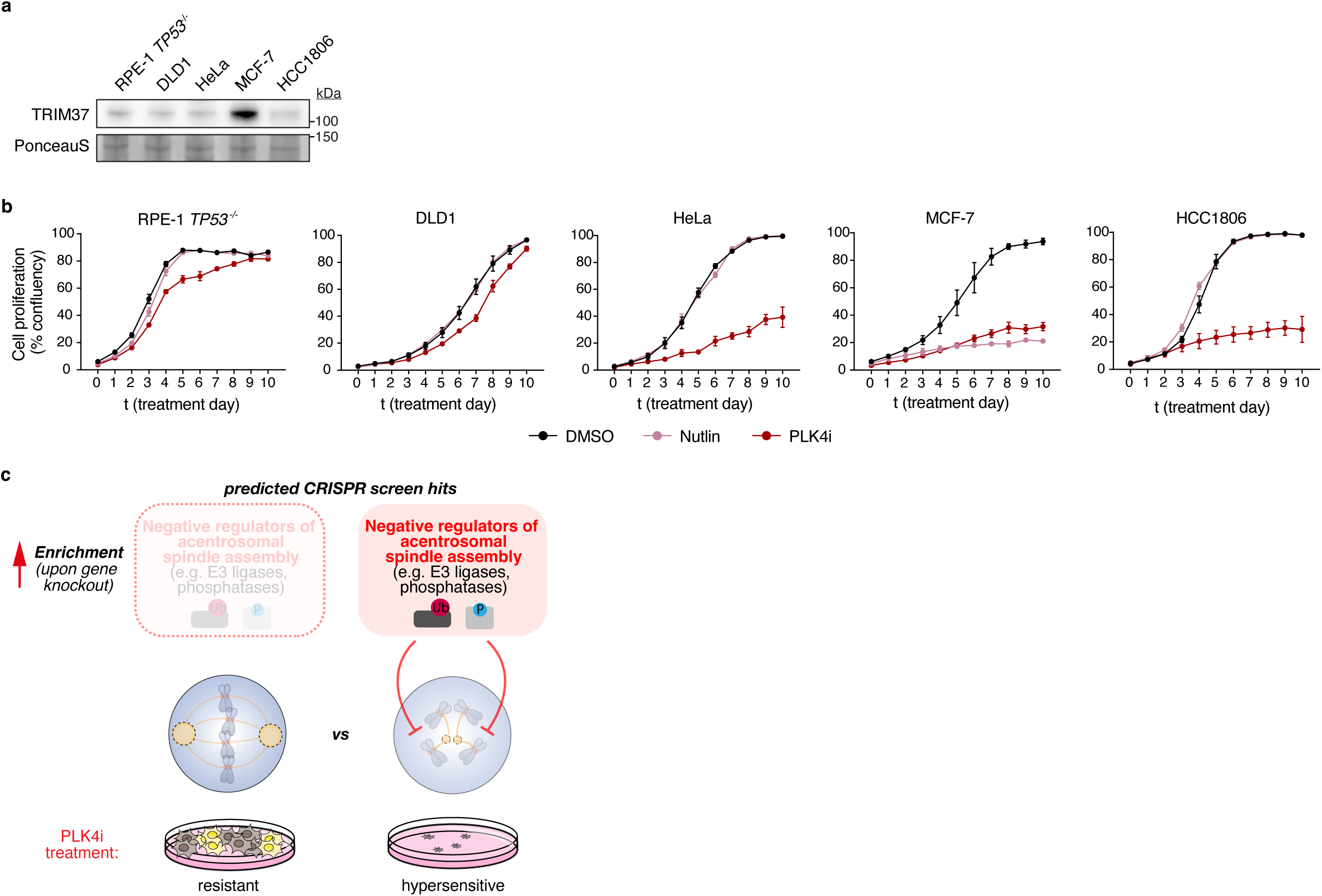
TRIM37 expression, PLK4i response across cancer cell lines, and predicted genetic contributors to PLK4i hypersensitivity. (related to Figure 1) **a**, Immunoblot showing relative TRIM37 expression levels in RPE1 *TP53^-/-^*, DLD1, HeLa, MCF-7 and HCC1806 cells. Ponceau-stained blot indicates loading. Representative data; *n* = 3 biological replicates. **b**, Cell proliferation curves of the cell lines shown in (a) treated with DMSO (control), PLK4i (500 nM), or the p53 activator Nutlin-3 (4 μM) for 10 d. *n* = 3 biological replicates. Mean ± s.e.m. **c**, Schematic illustrating the genome-wide CRISPR–Cas9 screening strategy and the predicted PLK4i-hypersensitivity suppressor hits. Negative regulators of acentrosomal spindle assembly that normally suppress spindle formation are predicted to be selectively enriched under PLK4i treatment, as their loss enables bipolar spindle assembly and confers resistance to PLK4 inhibition.

**Extended Data Figure 2.**
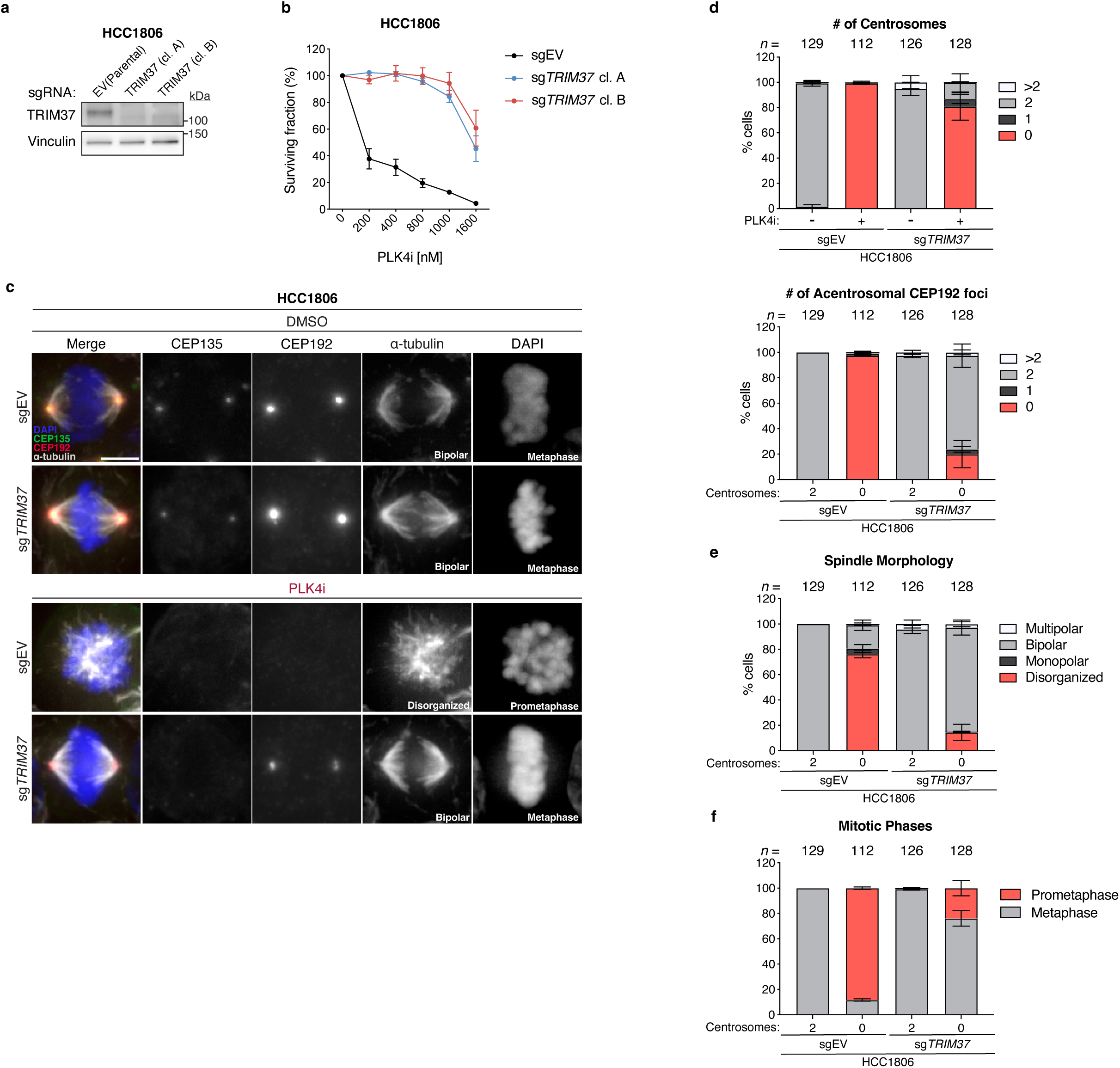
*TRIM37* loss suppresses PLK4i hypersensitivity and restores CEP192 foci at acentrosomal spindle poles in HCC1806 cells. **a**, Immunoblot showing TRIM37 levels in parental HCC1806 and two TRIM37 knockout clones. Vinculin, loading control. Representative data; *n* = 3 biological replicates. EV, empty vector. **b**, PLK4i sensitivity of the cell lines shown in (**a**), plotted as surviving fraction across increasing PLK4i concentrations and quantified by a resazurin-based viability assay. *n* = 3 biological replicates. Mean ± s.e.m. **c**, Representative immunofluorescence images of mitotic HCC1806 sgEV and sg*TRIM37* cells treated with DMSO (control) or PLK4i (500 nM) for 6 d. *n* = 3 biological replicates. Scale bar = 5 μm. **d**–**f**, Quantification of centrosome number and acentrosomal CEP192 foci (**d**), spindle morphology (**e**), and mitotic phase distribution (**f**) of HCC1806 sgEV and sg*TRIM37* cells under the treatment conditions shown in **c**. n = 3 biological replicates. Mean ± s.e.m.

**Extended Data Figure 3.**
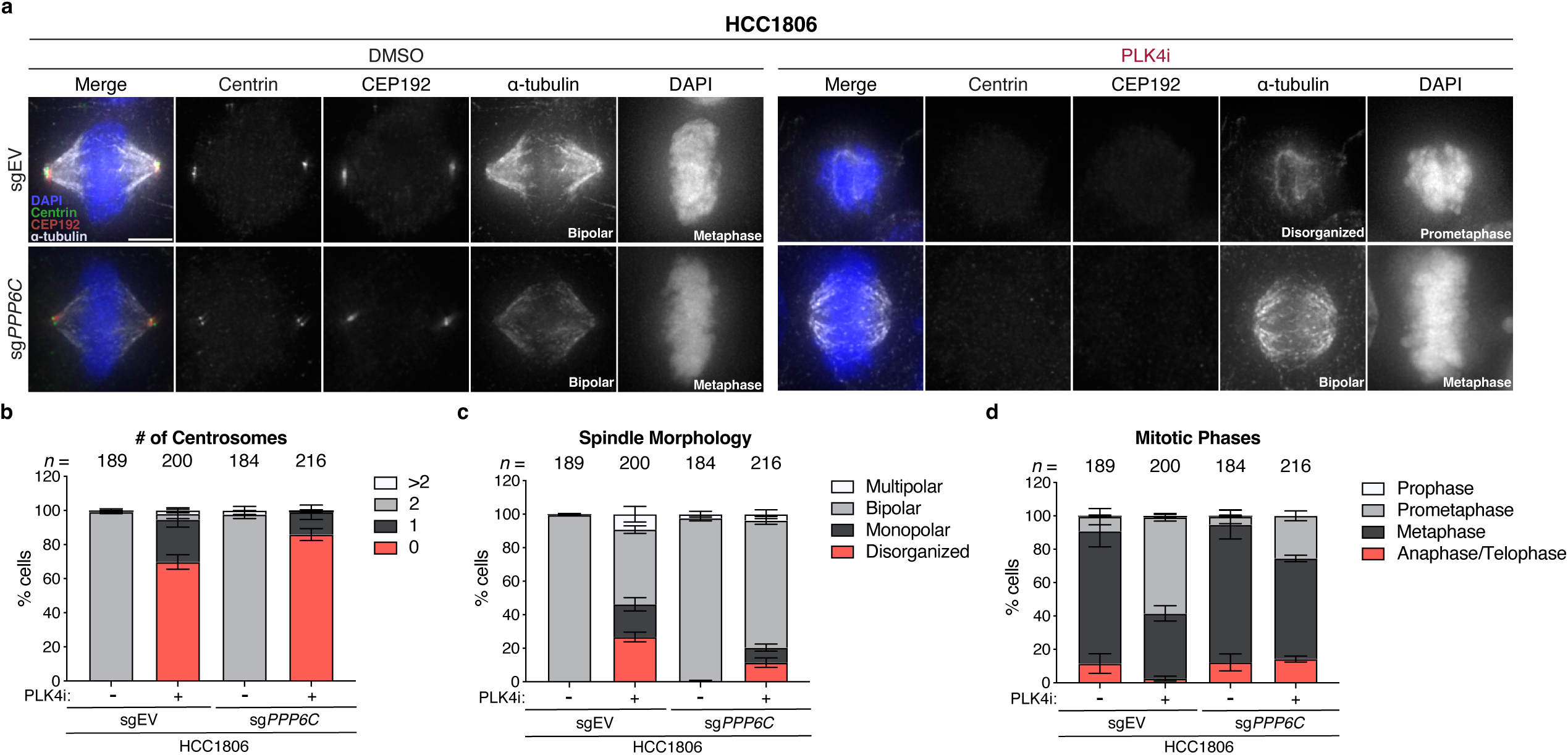
PPP6C loss restores spindle assembly under PLK4 inhibition independently of CEP192 foci at acentrosomal poles. (related to Figure 2) **a**, Representative immunofluorescence images of mitotic HCC1806 sgEV and sg*PPP6C* cells treated with DMSO (control) or PLK4i (500 nM) for 6 d. *n* = 3 biological replicates. Scale bar = 5 μm. **b**–**d**, Quantification of centrosome number (**b**), spindle morphology (**c**), and mitotic phase distribution (**d**) of HCC1806 sgEV and sg*PPP6C* cells under the treatment conditions shown in **c**. n = 3 biological replicates. Mean ± s.e.m.

**Extended Data Figure 4.**
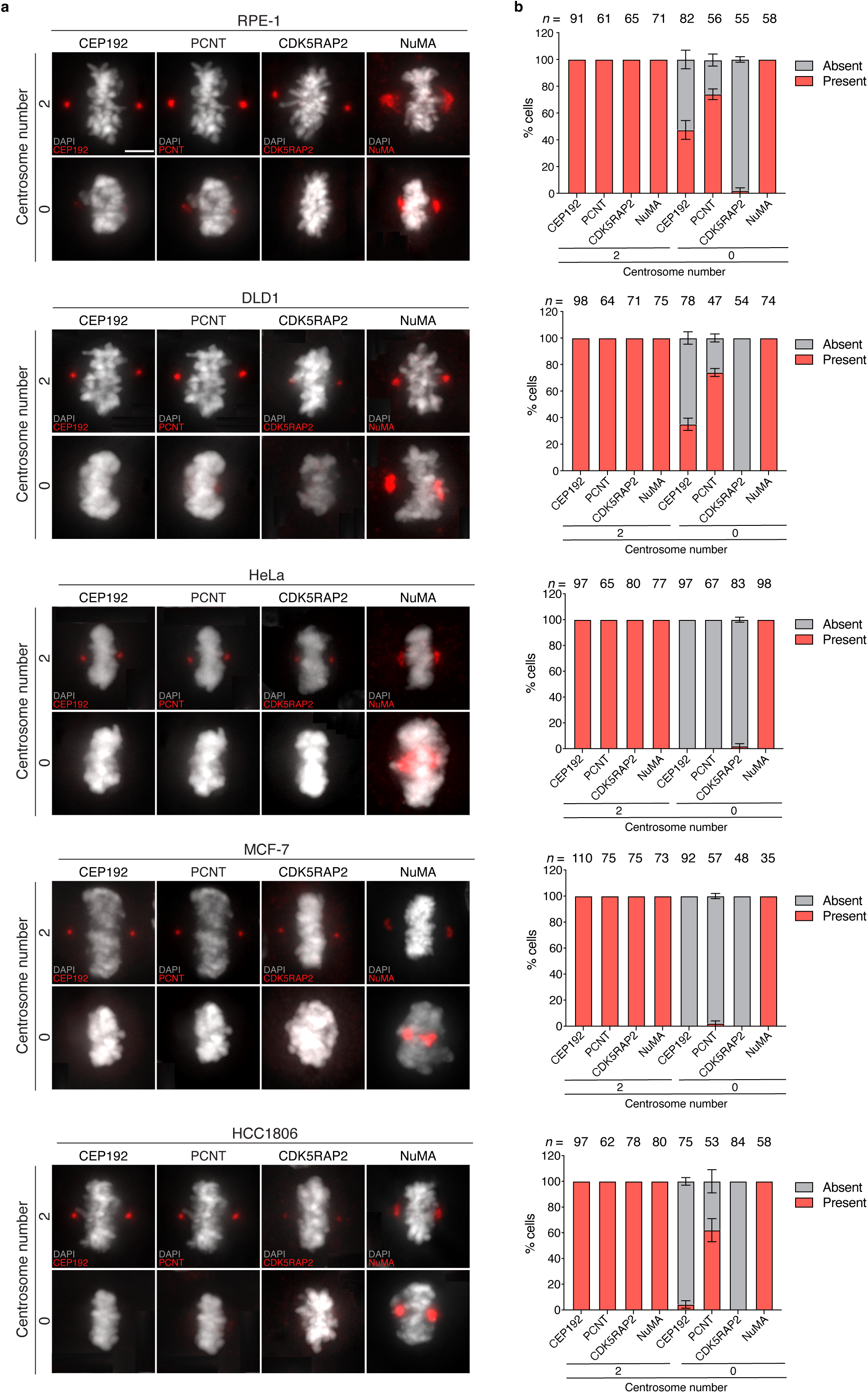
NuMA is a universal pole scaffold in acentrosomal spindles, whereas PCM retention varies across cell lines. (related to Figure 4) **a**, Representative immunofluorescence images of the indicated cell lines treated with DMSO (control) or PLK4i to induce centrosome depletion. Scale bar = 5 μm. n = 2 biological replicates. b, Quantification of the percentage of mitotic cells with detectable canonical PCM components (CEP192, PCNT, and CDK5RAP2) and NuMA at metaphase spindle poles under the conditions shown in (a). n = 2 biological replicates. Mean ± s.e.m.

**Extended Data Figure 5.**
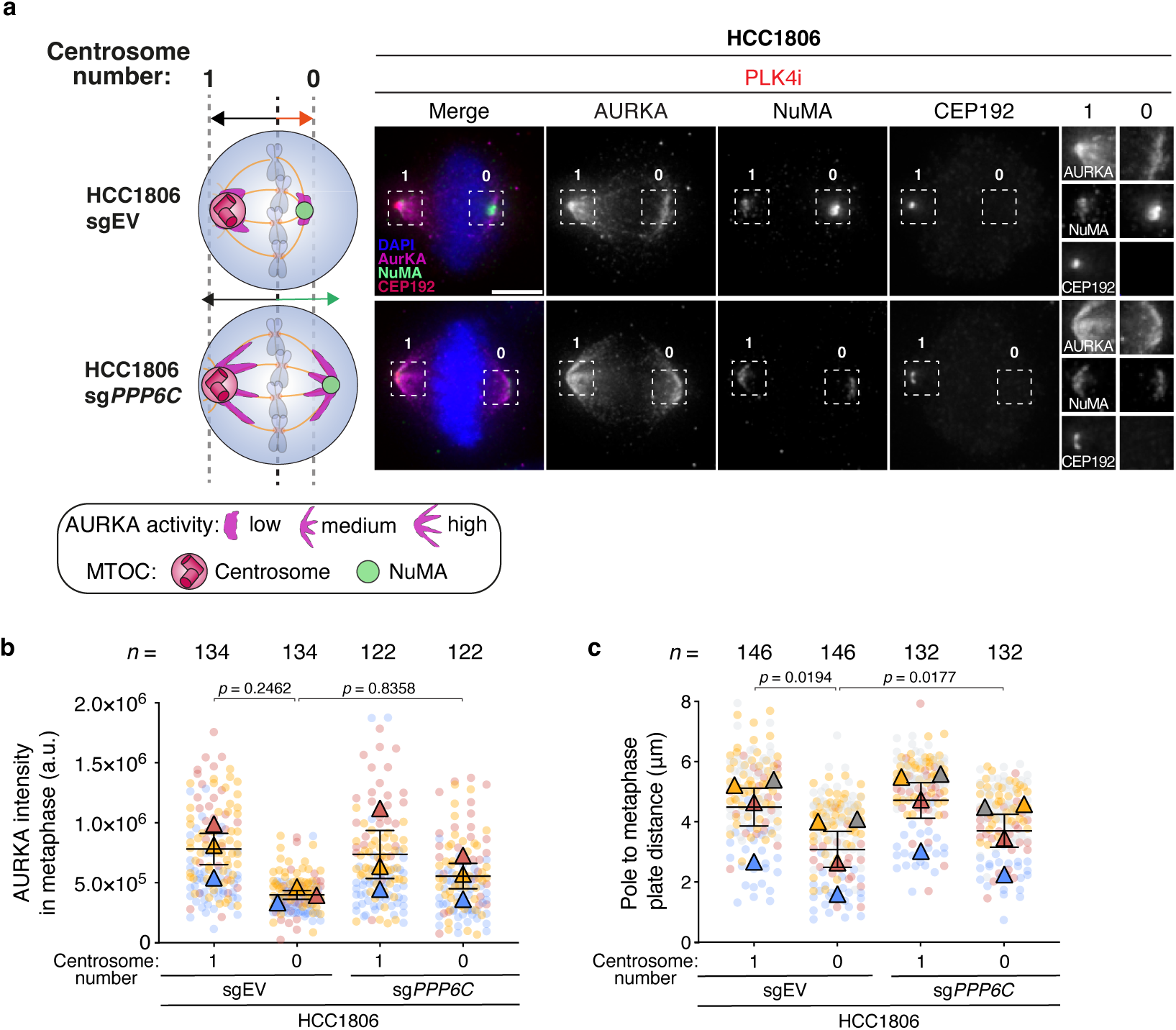
Reduced AURKA activity and shortened acentrosomal spindle pole length are partially offset by PPP6C loss. **a**, Left, Schematic illustrating the distinction between a centrosome-containing (1) pole (CEP192-positive) and an acentrosomal (0) pole (NuMA-positive, CEP192-negative), together with the corresponding AURKA localization patterns and half-spindle lengths. Right, Representative immunofluorescence images of mitotic HCC1806 sgEV and sg*PPP6C* cells subjected to short-term PLK4 inhibition to generate one-centrosome cells. Scale bar = 5 μm. *n* = 3 biological replicates. Mean ± s.e.m. **b**, Quantification of AURKA intensity at centrosome-containing and acentrosomal poles in sgEV and sg*PPP6C* cells shown in (a). *n* = 3 biological replicates. *P* values were determined using a one-way ANOVA with a post hoc Tukey’s multiple-comparisons test. Mean ± s.e.m. **c**, Quantification of pole-to-metaphase plate distance (half-spindle length) for centrosome-containing and acentrosomal poles in sgEV and sgPPP6C cells shown in (a). n = 3 biological replicates. *P* values were determined using a one-way ANOVA with a post hoc Tukey’s multiple-comparisons test. Mean ± s.e.m.

**Extended Data Figure 6.**
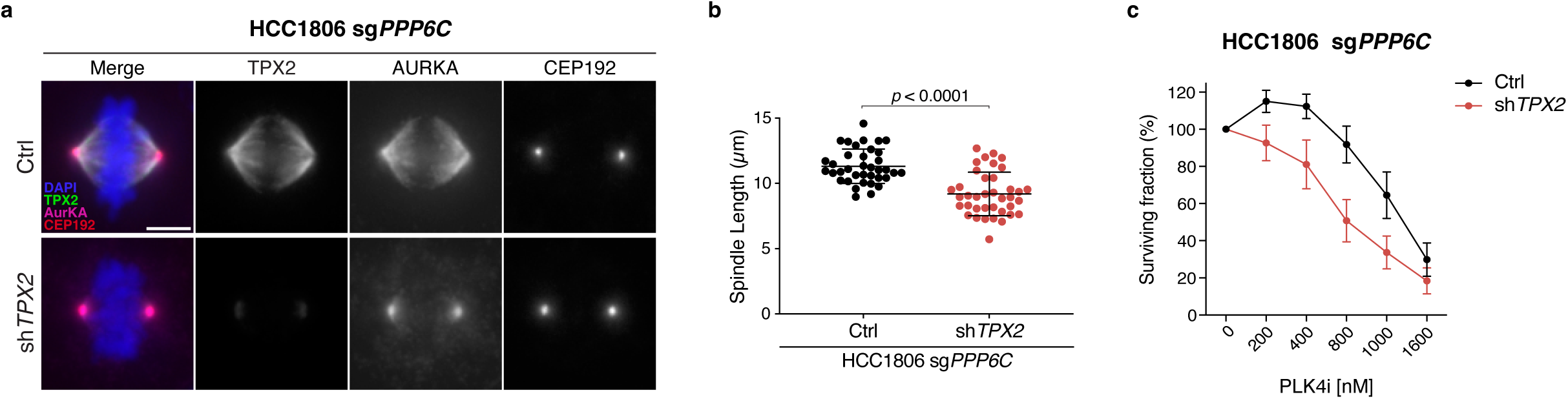
TPX2 is required to suppress PLK4i hypersensitivity in HCC1806 sg*PPP6C* cells. **a**, Representative immunofluorescence images of mitotic HCC1806 sg*PPP6C* cells transduced with vector control (shCtrl) or TPX2-targeting shRNA (shTPX2). Scale bar = 5 μm. *n* = 3 biological replicates. **b,** Quantification of spindle length in HCC1806 sg*PPP6C* cells shown in (a). *n* = 3 biological replicates. *P* values were determined using an unpaired two-tailed *t*-test. Mean ± s.e.m. **c**, PLK4i sensitivity of the cell lines shown in (**a**), plotted as surviving fraction across increasing PLK4i concentrations and quantified by a resazurin-based viability assay. *n* = 3 biological replicates. Mean ± s.e.m.

**Extended Data Figure 7.**
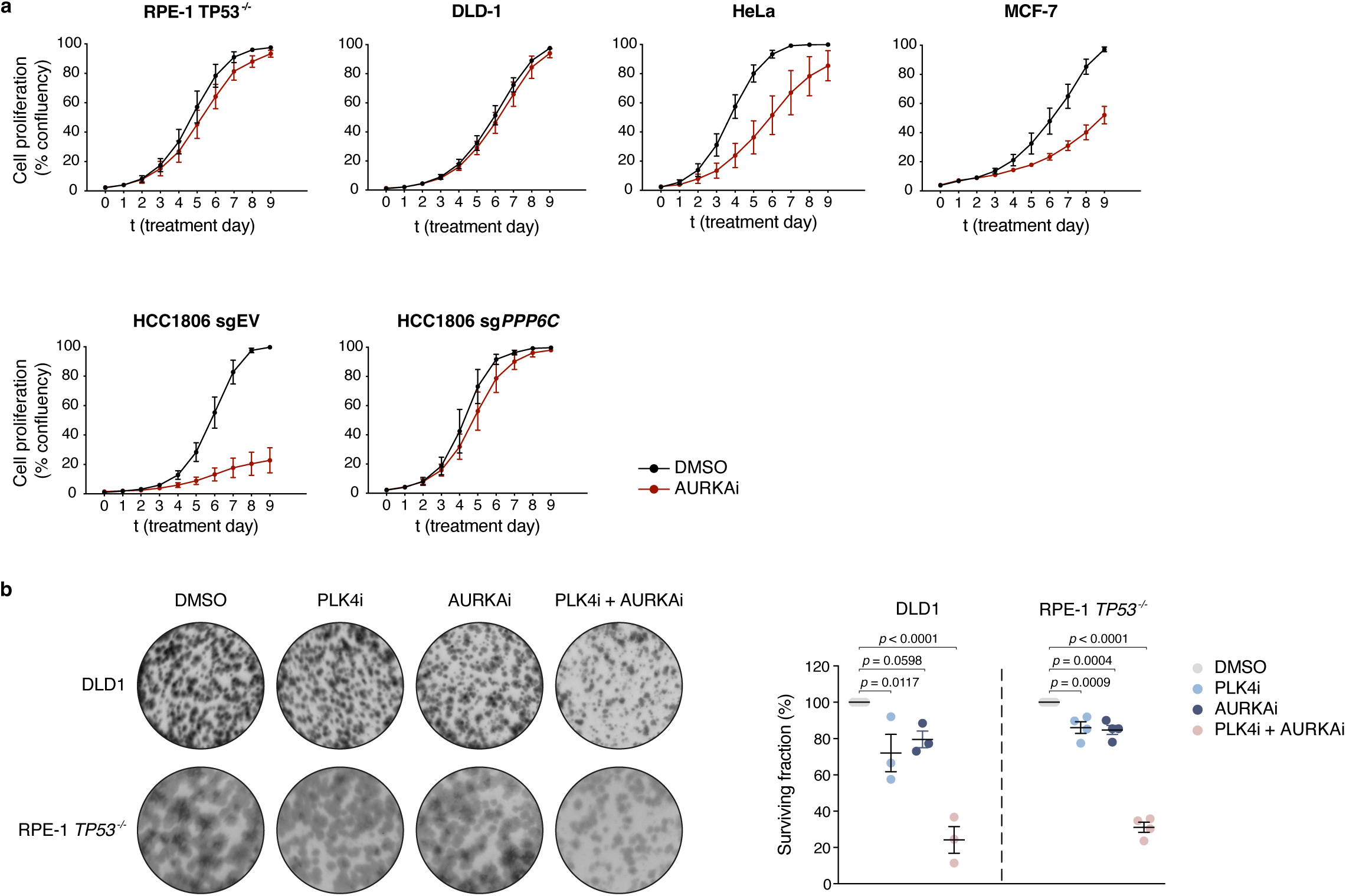
Intrinsic and inducible AURKA dependency in cancer cell lines. **a**, Cell proliferation curves of the indicated cell lines treated with DMSO (control) or the AURKA inhibitor MLN8237 (45 nM) for 9 d. *n* = 3 biological replicates. Mean ± s.e.m. **b**, Left, Representative 14-d clonogenic survival assay of the indicated cell lines treated with DMSO (control), PLK4i (500 nM), AURKAi (45 nM), or the combination of PLK4i and AURKAi. *n* = 3 biological replicates. Right, Quantification of surviving fraction. *P* values were determined using a one-way ANOVA with a post hoc Dunnett’s multiple-comparisons test to evaluate each treatment relative to DMSO control. Mean ± s.e.m.

## Methods

### Cell lines and culture conditions

hTERT *TP53*^−*/*−^ RPE-1, DLD1, HeLa and MCF-7 cells were grown in DMEM medium (Corning Cellgro) containing 10% fetal bovine serum (Sigma), 100 U/ml penicillin, 100 U/ml streptomycin and 2 mM L-glutamine. HCC1806 cells were grown in RPMI 1640 medium (Corning Cellgro) containing 10% fetal bovine serum (Sigma), 100 U/mL penicillin, 100 U/mL streptomycin and 2 mM L-glutamine. All cell lines were maintained at 37°C in a 5% CO_2_ atmosphere with 21% oxygen and routinely checked for mycoplasma contamination.

### Gene targeting and stable cell lines

To generate CRISPR/Cas9-edited lines, sgRNAs targeting *PPP6C* (5′-tgagagtagacagataacac-3′) and *TRIM37* (5′-ctccccaaagtgcacactga-3′) were cloned into the lentiCRISPR v2 vector (#52961; Addgene) modified for blasticidin resistance. Lentiviral particles were produced as described below. Cells were transduced and stable polyclonal populations of cells selected and maintained in the presence of 4 µg/mL blasticidin. Monoclonal cell lines were isolated by limiting dilution, and the ablation of protein production was assessed by immunoblotting.

To generate fluorescent histone H2B–iRFP- and EGFP–α-tubulin-expressing cell lines, the corresponding open reading frames (ORFs) were cloned into FUGW lentiviral vectors. Fluorescent HCC1806 cell populations were generated by lentivirus-mediated transduction and isolated by fluorescence-activated cell sorting (FACS).

### RNA interference

A Dharmacon pGIPZ lentiviral vector containing shRNA targeting TPX2 (5′-ttagcagtggaatcgagtg-3′) was purchased (Horizon). Stable shRNA-mediated knockdown (KD) cell lines were generated by lentivirus-mediated transduction. Polyclonal populations of cells were subsequently selected and maintained in the presence of puromycin (1.0 µg/mL). Knock down efficiency was assessed by immunoblotting.

Cells were transfected with Silencer Select siRNA targeting NuMA1 (Assay ID s9776; #4392420, Thermo Fisher Scientific) or non-targeting Silencer Select Negative Control #1 siRNA (#4390843, Thermo Fisher Scientific) using Lipofectamine RNAiMAX (Invitrogen) according to the manufacturer’s protocol, at a final siRNA concentration of 20 nM. Knockdown efficiency was assessed by immunofluorescence microscopy.

### Lentiviral production and transduction

Lentiviral expression vectors were co-transfected into 293FT cells with the lentiviral packaging plasmids psPAX2 and pMD2.G (Addgene #12260 and #12259). Briefly, 3 x 10^6^ 293FT cells were seeded into a Poly-L-Lysine coated 10 cm culture dish the day before transfection. For each 10 cm dish the following DNA were diluted in 0.6 mL of OptiMEM (Thermo Fisher Scientific): 4.5 µg of lentiviral vector, 6 µg of psPAX2 and 1.5 µg of pMD2.G. Separately, 72 µl of 1 µg/µl 25 kDa polyethyleneimine (PEI; Sigma) was diluted into 1.2 mL of OptiMEM, briefly vortexed, and incubated at room temperature for 5 min. After incubation, the DNA and PEI mixtures were combined, briefly vortexed, and incubated at room temperature for 20 min. During this incubation, the culture media was replaced with 17 mL of pre-warmed DMEM + 10% FBS. The transfection mixture was then added dropwise to the 10 cm dish. Viral particles were harvested 48 h after the media change and filtered through a 0.45 µm PVDF syringe filter. The filtered supernatant was either concentrated in 100 kDa Amicon Ultra Centrifugal Filter Units (Millipore) or used directly to infect cells. Aliquots were snap-frozen and stored at -80°C. For transduction, lentiviral particles were diluted in complete growth media supplemented with 10 µg/mL polybrene (Sigma) and added to cells.

### Chemical inhibitors

Centrinone (Tocris Bioscience) was dissolved in DMSO and used at a final concentration of 500 nM. MLN8237 (Selleck Chemicals) was dissolved in dimethyl sulfoxide (DMSO) and used at a final concentration of 45 nM. (−)-Nutlin-3 (Cayman Chemical) was dissolved in ethanol and used at a final concentration of 4 μM.

### CRISPR–Cas9 genome-wide screen

Pooled genome-wide CRISPR–Cas9 knockout screens in HCC1806 cells were performed as described previously. HCC1806 cells were infected with lentiCas9-Blast (#52962; Addgene). Positive selection of transduced cells was performed 2 days post-transfection with 5 µg/mL blasticidin. Monoclonal cell lines were isolated by limiting dilution and Cas9 expression was validated by immunoblotting.

The human Brunello CRISPR knockout sgRNA library was purchased from Addgene (a gift of David Root and John Doench; #73178) and plasmid DNA amplified according to the manufacturer’s instructions. To produce virus, the Brunello pooled plasmid library and the lentiviral packaging plasmids psPAX2 and pMD2.G were co-transfected into 40 × 15 cm culture dishes of HEK293FT cells. 6 × 106 HEK293FT cells were seeded into a poly-l-Lysine-coated 15 cm culture dish the day before transfection. For each 15 cm dish, the following DNA was diluted in 1.2 mL OptiMEM (Thermo Fisher Scientific): 9 µg lentiviral vector, 12 µg psPAX2, and 3 µg pMD2.G. Separately, 70 µl of 1 µg/µL 25-kD polyethylenimine (PEI) (Sigma) was diluted into 1.2 mL OptiMEM and incubated at room temperature for 5 min. After incubation, the DNA and PEI mixtures were combined and incubated at room temperature for 20 min. During this incubation, the culture media was replaced with 16 mL pre-warmed DMEM + 10% FBS. The transfection mixture was then added dropwise to the 15 cm dish. Viral particles were harvested at 24, 48, and 72 h after the media change. Media collected from 24, 48, and 72 h were pooled and filtered through a 0.45 µm PVDF syringe filter. The media was then concentrated using Amicon Ultra-15 Centrifugal Filter Unit with Ultracel-50 membrane (EMD Millipore Corporation cat# UFC905024). The virus was then frozen and stored at −80°C.

Cells were transduced with the Brunello library via spinfection as previously described^34^. To determine the optimal virus volumes for achieving an MOI ∼ 0.3, each new batch of virus was titered by spinfecting 3 × 10^6^ cells with several different volumes of virus. Briefly, 3 × 10^6^ cells per well were seeded into a 12 well plate in growth media supplemented with 10 µg/mL polybrene. Each well received a different titrated virus amount (between 5 and 50 µL) along with a no-transduction control. The plate was centrifuged at 1000 g for 2 h at 35°C. After the spin, media was aspirated, and fresh growth media was added. The following day, cells were counted, and each well was split into duplicate wells. One well received 3 µg/mL puromycin (Sigma) for 3 days. Cells were counted and the percent transduction was calculated as the cell count from the replicate with puromycin divided by the cell count from the replicate without puromycin multiplied by 100. The virus volume yielding a MOI closest to 0.3 was chosen for large-scale transductions.

For the pooled screens, a theoretical library coverage of ≥250 cells per sgRNA was maintained at every step. Cells were infected at MOI ∼ 0.3 and selected with puromycin at 3 μg/ml for 3 days. MOI was calculated using a control well infected in parallel following the procedure outlined above. Infected cells were expanded under puromycin selection for 6 or 7 days and subsequently seeded into 15 cm dishes a day prior to treatment. PLK4i (800 nM) was applied throughout the treatment period, and cell pellets were taken after 40 days at which point the screen was terminated. DMSO vehicle was used as the negative control.

Genomic DNA was isolated using the QIAamp Blood Maxi Kit (Qiagen) per manufacturer’s instructions. Genome-integrated sgRNA sequences for each sample were amplified and prepared for Illumina sequencing using a two-step PCR procedure as previously described^34^. For the first PCR, a region containing the sgRNA cassette was amplified using primers specific to the sgRNA-expression vector (lentiGuide-PCR1-F: 5’-aatggactatcatatgcttaccgtaacttgaaagtatttcg-3’; lentiGuide-PCR1-R: 5’-ctttagtttgtatgtctgttgctattatgtctactattctttcc-3’). The thermocycling parameters for the first PCR were as follows: 98 °C for 1 min, 20 cycles of (98 °C for 30 s, 65 °C for 30 s, 72 °C for 30 s), and 72°C for 1 min. The resulting amplicons for each sample were pooled and purified using AMPURE XP beads (Beckman Coulter) with a bead to sample ratio of 0.6x and 1.0x for double size selection to exclude primers and genomic DNA. Primers for the second PCR include Illumina adapter sequences, a variable length sequence to increase library complexity and a 8 bp barcodes for multiplexing of different biological samples (F2: 5’-aatgatacggcgaccaccgagatctacactctttccctacacgacgctcttccgatct-[4–7 bp random nucleotides]-[8 bp barcode]-tcttgtggaaaggacgaaacaccg-3’; R2: 5’-caagcagaagacggcatacgagatgtgactggagttcagacgtgtgctcttccgatcttctactattctttcccctgcactgt-3’). 2.5 µL of the product from the first PCR reaction was used, and the thermocycling parameters for the second PCR were as follows: 98°C for 30 s, 12 cycles of (98°C for 1 s, 70°C for 5 s, 72°C for 35 s). Second PCR products were pooled, purified using AMPURE XP beads with a bead to sample ratio of 1.8x and quantified using the Qubit dsDNA BR Assay Kit (Thermo Fischer Scientific). Diluted libraries with 5% PhiX were sequenced with MiSeq (Illumina).

Sequencing data were processed for sgRNA representation using custom scripts. Briefly, sequencing reads were first demultiplexed using the barcodes in the forward primer and then trimmed to leave only the 20 bp sgRNA sequences. The spacer sequences were then mapped to the spacers of the designed sgRNA library using Bowtie^35^. For mapping, a maximum of one mismatch was allowed in the 20 bp sgRNA sequence. Mapped sgRNA sequences were then quantified by counting the total number of reads. The total numbers of reads for all sgRNAs in each sample were normalized.

We used the MaGeCK scoring algorithm (model-based analysis of genome-wide CRISPR-Cas9 knockout) to analyze and the rank the genes from the screens ^36^. β-scores and FDR values were also derived for each gene between DMSO or PLK4i conditions. Genes with a β-score outside 1.5 standard deviations from the population mean and with an FDR cutoff of ≤ 0.3 were taken forward for further validation. Graphing and downstream analysis was performed in R.

### Antibody techniques

For immunoblot analyses, protein samples were resolved by SDS-PAGE on pre-cast NuPAGE™ gels (Invitrogen) with molecular weight ladders (PageRuler Plus, Invitrogen). Following electrophoresis, proteins were transferred onto nitrocellulose membranes using a Mini Trans-Blot Cell (BioRad) wet transfer system and subsequently probed with the following primary antibodies: PPP6C (rabbit, Bethyl, A300-844A, 1:2000), TRIM37 (rabbit, Bethyl, A301-174A, 1:1000), vinculin (mouse, Santa Cruz Biotechnology, sc-73614, 1:1000), with SuperSignal West Pico PLUS or Femto Maximum chemiluminescence substrate (Thermo Fisher Scientific). Signals were visualized and acquired using a Genesys G:Box Chemi-XX6 system (Syngene).

For immunofluorescence, cells were cultured on 12-mm glass coverslips and fixed for 8 min in 100% ice-cold methanol at -20°C. Cells were blocked in 2.5% FBS, 200 mM glycine, and 0.1% Triton X-100 in PBS for 1 h. Antibody incubations were conducted in the blocking solution for 1 h. DNA was stained with DAPI, and cells were mounted in ProLong Gold Antifade (Invitrogen). Staining was performed with the following primary antibodies: NuMA (rabbit, Cell Signaling, #3888, 1:200), Aurora A (rabbit, Cell Signaling, #4718, 1:100), Aurora A (mouse, Abcam, ab13824, 1:200), Phospho-Aurora A (Thr288) (rabbit, Cell Signaling, #3079, 1:1600), CDK5RAP2 (rabbit, Millipore, 06-1398, 1:1000), PCNT (rabbit, Abcam, ab4448, 1:1000), TPX2 (mouse, Abcam, ab32795, 1:500), CEP135-Alexa 555 (directly-labelled rabbit, homemade, 1:1000), CEP192-Cy5 (directly-labelled goat, raised against CEP192 residues 1–211, home-made, 1:1000), Centrin (mouse, Millipore, 04-1624, 1:1000), and α-tubulin (rat, Invitrogen, MA1-80017, 1:1000).

Immunofluorescence images were acquired using a DeltaVision Elite system (GE Healthcare) controlling a Scientific CMOS camera (pco.edge 5.5). Acquisition parameters were controlled by SoftWoRx suite (GE Healthcare). Images were collected at room temperature (25°C) using an Olympus 40x 1.35 NA, 60x 1.42 NA or Olympus 100x 1.4 NA oil objective at 0.2 μm z-sections. Images were acquired using Applied Precision immersion oil (N=1.516). For quantitation of signal intensity at the centrosome, deconvolved 2D maximum intensity projections were saved as 16-bit TIFF images. Signal intensity was determined using ImageJ by drawing a circular region of interest (ROI) around the centriole (ROI S). A larger concentric circle (ROI L) was drawn around ROI S. ROI S and L were applied to the channel of interest and the signal in ROI S was calculated using the formula IS − [(IL − IS/AL − AS) × AS], where A is area and I is integrated pixel intensity. Quantification of spindle length was performed using the ImageJ straight-line tool by drawing a line from one pole to the other.

### Live-cell microscopy

Fluorescent cell lines were seeded into µ-Slide 4-well or 8-well glass bottom chamber slides (Ibidi) and treated with the indicated drugs 24 h later. Time-lapse imaging was performed 72 h post-drugging using a Zeiss Axio Observer 7 inverted microscope equipped with Slidebook 2024 software (3i—Intelligent, Imaging Innovations, Inc.), X-Cite NOVEM-L LED laser and filter cubes, and a Prime 95B CMOS camera (Teledyne Photometrics) with a 40×/1.3 plan-apochromat oil immersion objective. During imaging, cell conditions were maintained at 37°C, with 5% CO_2_, and 60% relative humidity (RH) using a stage top incubator (Okolab).

For analysis of mitotic phenotypes, cells were treated as indicated and imaged every 5 min in 9 × 3 μm z-sections in the respective fluorescent channels. Phenotypic evaluation and intensity measurements were performed on maximum intensity projected 2D time-lapse images. Spindle status was assessed from nuclear envelope breakdown to mitotic exit, based on the number, position, and relative distance of spindle poles determined through qualitative observation. These observations were standardized using a set of pre-determined reference images.

### Cell proliferation and viability assays

Cells seeded in triplicate in 6-well plates were treated with the indicated drugs 16 h later. After the specified number of days, cells were fixed and stained using 0.5% (w/v) crystal violet in 20% (v/v) methanol for 5 minutes. Excess stain was rinsed thoroughly with distilled water, and plates were air-dried overnight. For quantification, bound crystal violet was dissolved in 10% (v/v) acetic acid in dH_2_O and absorbance of 1:50 dilutions were measured at 595 nm using a Synergy HT Microplate Reader (BioTek Instruments). The optical density at 595 nm (OD_595_) served as a quantitative metric of relative cell growth.

For longitudinal cell proliferation curve measurements, cells were seeded in the 96-well plates. The following day, drugs were added, and confluency measurements were taken every 4 h for 7 days using the CellCyte X live-cell imaging system (Cytena). To assay for surviving fraction, the culture medium was replaced with phenol red-free DMEM (Thermo Fisher Scientific) containing 10% fetal bovine serum (Sigma), 100 U/ml penicillin, 100 U/ml streptomycin, 2 mM L-glutamine, and 10 μg/mL resazurin (Sigma). Plates were incubated for 2–4 h, or until the medium in untreated control wells developed a pink colour. Relative fluorescence was measured using a Synergy HT microplate reader (BioTek Instruments) at 530 nm (excitation) and 590 nm (emission)

### Statistical analysis

GraphPad Prism (version 10) was used for graphing and statistical analyses. Specific statistical methods for each experiment are provided in the figure legends.

## Data availability

Data that support the findings of this study are available from the corresponding authors upon reasonable request. Source data are provided with the paper.

## Supplementary Information

**Supplementary Video 1.**

DMSO-treated sgEV HCC1806 cell undergoing normal mitotic progression, as shown in Fig. 2d. H2B–iRFP is shown in red and TagRFP–tubulin in gray. Images were acquired as 3-μm z-sections, with one stack collected every 5 min and displayed at 8 frames per second.

**Supplementary Video 2.**

PLK4i-treated sgEV HCC1806 cell displaying unstable bipolar spindle organization and prolonged mitosis culminating in mitotic death, as shown in Fig. 2d. H2B–iRFP is shown in red and TagRFP–tubulin in gray. Images were acquired as 3-μm z-sections, with one stack collected every 5 min and displayed at 8 frames per second.

**Supplementary Video 3.**

DMSO-treated sg*PPP6C* HCC1806 cell undergoing normal mitotic progression, as shown in Fig. 2d. H2B–iRFP is shown in red and TagRFP–tubulin in gray. Images were acquired as 3-μm z-sections, with one stack collected every 5 min and displayed at 8 frames per second.

**Supplementary Video 4.**

PLK4i-treated sg*PPP6C* HCC1806 cell maintaining stable bipolar spindle organization and exiting mitosis normally, as shown in Fig. 2d. H2B–iRFP is shown in red and TagRFP–tubulin in gray. Images were acquired as 3-μm z-sections, with one stack collected every 5 min and displayed at 8 frames per second.

